# Ninjurin2 Regulates Vascular Smooth Muscle Cell Phenotypic Switching and Vascular Remodeling through Interacting with PDGF Receptor-β

**DOI:** 10.1101/2025.05.27.656052

**Authors:** Mengru Jia, Jiahui Zhuo, Xinming Zhao, Ting Liu, Huifang Zhang, Kexin Xu, Pengyun Wang, Chengqi Xu

## Abstract

Vascular smooth muscle cell (VSMC) phenotypic switching from a contractile to synthetic state plays a pivotal role in vascular remodeling. Although genome-wide association studies have linked *NINJ2* (encode Ninjurin2) polymorphisms are associated with risk of atherosclerosis and related conditions like ischemic stroke and coronary artery disease, its functional role in regulating VSMC plasticity remains unexplored. Here, using Ninj2 knockout (*Ninj2^-/-^*) and smooth muscle-specific overexpressing (*Ninj2^smcTG^*) mice subjected to carotid artery ligation, we demonstrate that Ninjurin2 deficiency exacerbates injury-induced intimal hyperplasia, whereas its overexpression attenuates this pathological remodeling. Mechanistically, Ninjurin2 knockdown promoted PDGF-BB-mediated VSMC phenotypic switching, proliferation, and migration, while biochemical studies revealed that Ninjurin2 interacts with PDGFRβ to suppress FAK signaling. These findings identify Ninjurin2 as a novel regulator of VSMC phenotypic switching via the PDGFRβ/FAK axis, offering new mechanistic insights into vascular remodeling and potential therapeutic avenues for atherosclerosis.

## 1. Introduction

Cardiovascular disease (CVD) is the leading cause of morbidity and mortality globally. Vascular smooth muscle cells (VSMCs), the predominant cell type in blood vessels, play a pivotal role in both sustaining vascular homeostasis and contributing to vascular remodeling[1]. VSMCs are typically quiescent and provide the contractile force necessary for blood vessels to regulate vascular function[2]. Upon vascular injury, VSMCs undergo phenotypic switching from a contractile to a synthetic state, characterized by increased proliferation and migration, which significantly contributes to intimal hyperplasia and vascular remodeling observed in numerous cardiovascular diseases[3, 4].

In a previous genome-wide association study (GWAS), two single nucleotide polymorphisms (SNPs), rs11833579 and rs12425791, located near the *NINJ2* (encode Ninjurin2) gene on chromosome 12p13, were significantly correlated with the risk of ischemic stroke[5]. Additionally, another SNP, rs34166160, was identified as associated with ischemic stroke risk within the Cohorts for Heart and Aging in Genomic Epidemiology (CHARGE) consortium and with coronary artery disease (CAD) in a Chinese population[6, 7]. In a separate Chinese cohort, rs12425791 and rs11833579 were identified as independent predictors of stroke-related mortality or stroke recurrence in patients with incident ischemic stroke[8]. Given that atherosclerosis is the primary pathological underpinning of both ischemic stroke and coronary artery disease, the *NINJ2* gene is a plausible candidate gene that may play a critical role in atherosclerotic cardiovascular diseases and vascular remodeling. However, the specific mechanisms by which *NINJ2* influences ischemic stroke or atherosclerosis remain to be elucidated.

In the aorta, Ninjurin2 is expressed in both vascular endothelial cells and VSMCs[9, 10]. *NINJ2* activates endothelial cells via the Toll-like receptor 4 (TLR4) signaling pathway, promoting the adhesion of monocytes and macrophages[9]. Paradoxically, despite its expression in VSMCs, Ninjurin2’s role in regulating VSMC plasticity remains unknown. Intriguingly, a previous study demonstrated that in glioma cells, Ninjurin2 co-immunoprecipitates with platelet-derived growth factor receptor-beta (PDGFR-β)[11], which is crucial for the phenotypic switching of VSMCs[12]. PDGFR-β, activated by its natural ligand PDGF-BB—a potent mitogen and chemoattractant for VSMCs—initiates downstream signaling pathways that contribute to the phenotypic switching of VSMCs, vascular remodeling, and the development of various vascular diseases[13, 14]. Based on these findings, it is plausible to hypothesize that Ninjurin2 may regulate the phenotypic switching of VSMCs and vascular remodeling, thereby influencing the pathogenesis of atherosclerotic cardiovascular diseases.

To elucidate the role of *NINJ2* in vascular remodeling, the current study investigates *NINJ2*’s involvement in vascular injury and remodeling. We employed carotid artery ligation injury mouse models, which included *Ninj2* knockout mice and mice with VSMC-specific *Ninj2* overexpression, complemented by in vitro experiments. Subsequently, we explored the potential mechanisms by which Ninjurin2 regulates the phenotypic switching of VSMCs and vascular remodeling.

## 2. Materials and Methods

### 2.1 Mouse models

*Ninj2* gene knockout mice (*Ninj2*^-/-^) were generated using TALEN technology, as previously described[15]. A deletion of an A base in exon 2 of *Ninj2* gene in C57BL/6 mice resulted in a frameshift mutation, leading to premature termination of protein translation. Mutations in the *Ninj2* gene were confirmed by DNA sequencing. Genotyping of the *Ninj2* knockout mice was performed using PCR with the following primers: forward (5′-CCAAGAGGACATTTCATCGCCTT-3′); and reverse (5′-TGATTTCAGCCGCATGGCATT-3′).

A knock-in conditional transgenic mouse line, *Rosa26-CAG-Loxp-stoploxp-Ninj2-IRES-EGFP* (*Ninj2^fl/fl^* mice), was constructed using CRISPR/Cas9-mediated gene knock-in technology as previously described[16]. The line targets *Ninj2-IRES-EGFP* into the *ROSA26* locus, with a loxP-flanked stop sequence preceding it. To achieve VSMC specific *Ninj2* overexpression, *Ninj2^fl/fl^* mice then crossed with *Tagln*-Cre transgenic mouse line, resulting in VSMC specific *Ninj2* overexpression mice (*Ninj2*^smcTG^).

Animal care and procedures were reviewed and approved by the Institutional Committee on Animal Care and Use, School of Medicine, Huazhong University of Science and Technology (No. [2016] ICUC469), and were conducted in compliance with institutional and relevant guidelines. The mice were housed in a specific pathogen-free (SPF) facility, maintained at a controlled temperature of 22±2℃, humidity of 55%±5%, and a 12-hour light/dark cycle, and provided with *ad libitum* access to a standard chow diet (WQJX BIO-TECHNOLOGY). Animals were anesthetized via intraperitoneal injection of pentobarbital sodium at a dosage of 50mg/kg body weight.

### 2.2 Mouse carotid artery ligation model

Carotid artery ligation (CAL) was performed on 10-week-old male *Ninj2*^-/-^ mice, wild-type (WT) littermates, *Ninj2*^smcTG^ and *Ninj2^fl/fl^* control mice according to the protocol established by Kumar et al[17]. The left common carotid artery was exposed and completely ligated just distal to the carotid bifurcation. The right common carotid artery was exposed as a non-ligated control (sham group). Post-operative analgesia was administered with buprenorphine at a dosage of 0.05 mg/kg every 12 hours for 48 hours. After a 28-day postoperative period, the mice were euthanized by cervical dislocation, and the arteries with the ligature knots were harvested for hematoxylin and eosin (H&E) staining to assess neointima formation[18].

### 2.3 Histological experiment

For histological analysis, the ligatured arteries were fixed in 4% paraformaldehyde at 4°C for 16 hours prior to paraffin embedding. Serial sections (4 μm) were obtained and stained with hematoxylin and eosin as previously described. For immunohistochemical staining, dewaxing of blood vessel sections was followed by antigen repair for 30 min. The slices were then combined with anti-GFP primary antibody (19245S; 1:200; CST) at 4 °C overnight, followed by biotin secondary antibody for 30 min. DAB substrate was used to stain the slides and hematoxylin was used for final detection.

### 2.4 Cell culture and siRNA transfection

Mouse aortic smooth muscle cells (MOVAS), HeLa, and HEK 293T cells were cultured in Dulbecco’s Modified Eagle Medium (DMEM) supplemented with 10% fetal bovine serum (FBS) and 1% penicillin-streptomycin solution (100×). Cultures were maintained at 37°C in a 5% CO2 incubator. The siRNA sequence of mouse *Ninj2* is 5 ‘-TGTTCATCGCCATCCTGAATT-3’ and the negative control si-NC sequence is 5 ‘-TTCTCCGAACGTGTCACGTTT-3’. Transfection was performed using Lipofectamine 2000 reagent (Invitrogen) in serum-free medium.

### 2.5 Quantitative reverse-transcription polymerase chain reaction (qRT–PCR)

cDNA was synthesized using the reverse transcription kit (Vazyme; R223-01). Quantitative real-time polymerase chain reaction (qPCR) analysis was conducted using the FastStart Universal SYBR Green Master (Roche) on a QuantStudio 6 Flex Real-Time PCR System (Applied Biosystems), following the manufacturer’s protocol. Primer sequences are provided in the supplementary table. Gene expression levels were normalized to TATA-box binding protein (TBP) and calculated using the 2*-^ΔΔCt^* method.

### 2.6 Western Blot

Tissue and cell samples were lysed using IP lysates supplemented with protease and phosphatase inhibitors (MedChem Express; HY-K0010; HY-K0021). Subsequently, 5× protein loading buffer was added for storage. Proteins were electrophoretically separated by SDS-PAGE and transferred to a PVDF membrane. The membrane was blocked with 5% skim milk powder, followed by an overnight incubation with the primary antibody. On the subsequent day, the samples were incubated with corresponding secondary antibodies, and protein bands were visualized using a Bio-Rad Imager. The antibodies used were as follows: GAPDH (60004-1-Ig; 1:1000 dilution; Proteintech), HSP90 (13171-1-AP, 1:1000 dilution; Proteintech), PDGFRβ (13449-1-AP; 1:1000 dilution; Proteintech), α-SMA (14395-1-AP; 1:1000 dilution; Proteintech), SM22 (10493-1-AP; 1:1000 dilution; Proteintech), FAK (112636-1-AP; 1:1000 dilution; Proteintech), AKT (6022033-2-Ig; 1:1000 dilution; Proteintech), Flag (66008-4-Ig; 1:1000 dilution; Proteintech), β-TUBULIN (10068-1-AP; 1:1000 dilution; Proteintech), pFAK (8556S, 1:1000 dilution, Cell Signaling Technology), pAKT (4060S, 1:1000 dilution, Cell Signaling Technology), GFP (598, 1:1000 dilution; MBL). Quantitative analysis of protein expression was performed using ImageJ software, with GAPDH/HSP90 serving as the internal reference proteins.

### 2.7 VSMC proliferation assay

MOVAS VSMCs were seeded in 12-well plates and cultured for 36 hours. Proliferation of VSMCs was quantitatively assessed using the BeyoClick™ EdU-488 Cell Proliferation Assay Kit (C0071S; Beyotime), following the manufacturer’s protocols. The assay measures VSMCs proliferation by detecting the incorporation of the nucleoside analog 5-ethynyl-2’-deoxyuridine (EdU) into newly synthesized DNA. This is achieved through a copper-catalyzed "click" reaction with a fluorescent dye, followed by fluorescence microscopy for photographic analysis of the samples.

### 2.8 VSMC migration assay

MOVAS-1 cells were transfected with *si-Ninj2* at a concentration of 50 nM and subsequently starved in a medium containing 0.5% FBS for 48 hours. A mechanical scratch was introduced using a pipette tip at the well bottom, followed by stimulation with recombinant human PDGF-BB (HY-P7087; MedChem Express) at a final concentration of 20 ng/mL. The cells were then allowed to migrate for 24 hours. Images were obtained using a Nikon ECLIPSE Ti microscope and analyzed with Image-Pro Plus 6.0 software. Cell migration was quantified as the ratio of (cell-free area at 0 hours - cell-free area at 24 hours) to the cell-free area at 0 hours.

Additionally, cell migration was assessed using Transwell assays in Boyden chambers. MOVAS cells were transfected with *si-Ninj2* (50 nM) for the Transwell migration experiment. After a 48-hour culture period, the cells were trypsinized to form a single-cell suspension. The lower chamber was filled with medium containing 20% FBS to induce migration, while 5000 cells were placed in the upper chamber. Following a 24-hour incubation, the chamber was stained with crystal violet and imaged using an inverted fluorescence microscope.

### 2.9 Immunofluorescence

Flag-*PDGFRB* and GFP-*NINJ2* plasmids were transfected into Hela cells and fixed with paraformaldehyde for 30 min after 48 h. After washing, 0.5% Triton X-100 was added. Flag polyclonal primary antibody(66008-4-Ig; 1:1000 dilution; Proteintech) was added for overnight incubation at 4 ℃ and Flag-PDGFRB protein was immunostained. After washing with PBS, the cells were incubated with secondary antibody at room temperature for 2 h. Hoechst 33342 stained the cell nuclei. Images were then taken with the FV3000 confocal microscope.

### 2.10 Immunoprecipitation Experiment

Flag-*PDGFRB* and GFP-*NINJ2* plasmids were transfected into HEK 293T cells and lysed after 48 h. The supernatant of cell lysate was collected, and 100 μL lysate was taken as Input positive control, and 5× protein loading buffer was added. The sample was incubated at 100 ℃ for 10 min and stored at -20 ℃. 500 μL was the IP group, incubated with Flag magnetic beads overnight; 400 μL was used as IgG negative control. IgG antibodies were added to agarose beads and incubated overnight. After washing with elution Buffer, add 1× protein loading buffer to cook sample for 10 min and store at -20 ℃.

### 2.11 Data Statistics and Analysis

All statistical analyses were performed by using GraphPad Prism 8 software, and data were presented as mean ±SEM. Statistical differences between two groups were analyzed using the Student’s t-test. Statistical differences among multiple groups were analyzed by one-way ANOVA followed by a Tukey post-hoc test. P-value < 0.05 was considered statistically significant. * P < 0.05, ** P < 0.01, ns, not significant.

## 3. Results

### 3.1. Ninj2 modulate neointimal formation and phenotypic switching of VSMC after carotid artery ligation in mouse

In order to elucidate the effect of Ninjurin2 on neointimal formation in vivo, we initially employed a mouse carotid artery ligation model in *Ninj2^-/-^*KO mice and wildtype (WT) littermates at the age of 10 weeks. After 28 days of carotid artery ligation, histological examination via hematoxylin and eosin (H&E) staining revealed a pronounced increase in neointimal formation, characterized by elevated intima-to-media ratios, in *Ninj2^-/-^*KO mice as compare with WT controls (Figure 1A-B). Moreover, we examined the VSMCs phenotypic switching between a contractile/differentiated state and a synthetic/dedifferentiated state by assessing the expression of contractile protein markers, including α-smooth muscle actin (α-SMA) and SM22 (transgelin). Western blot analysis demonstrated that, 28 days post-ligation, the expression of SM22 and α-SMA was significantly reduced in the ligatured arterial tissues of *Ninj2^-/-^*KO mice compare to WT controls (Figure 1C-E). Collectively, these findings imply that Ninjurin2 ablation exacerbates neointimal formation and modulates VSMC phenotypic switching following vascular injury.

**Figure 1.**
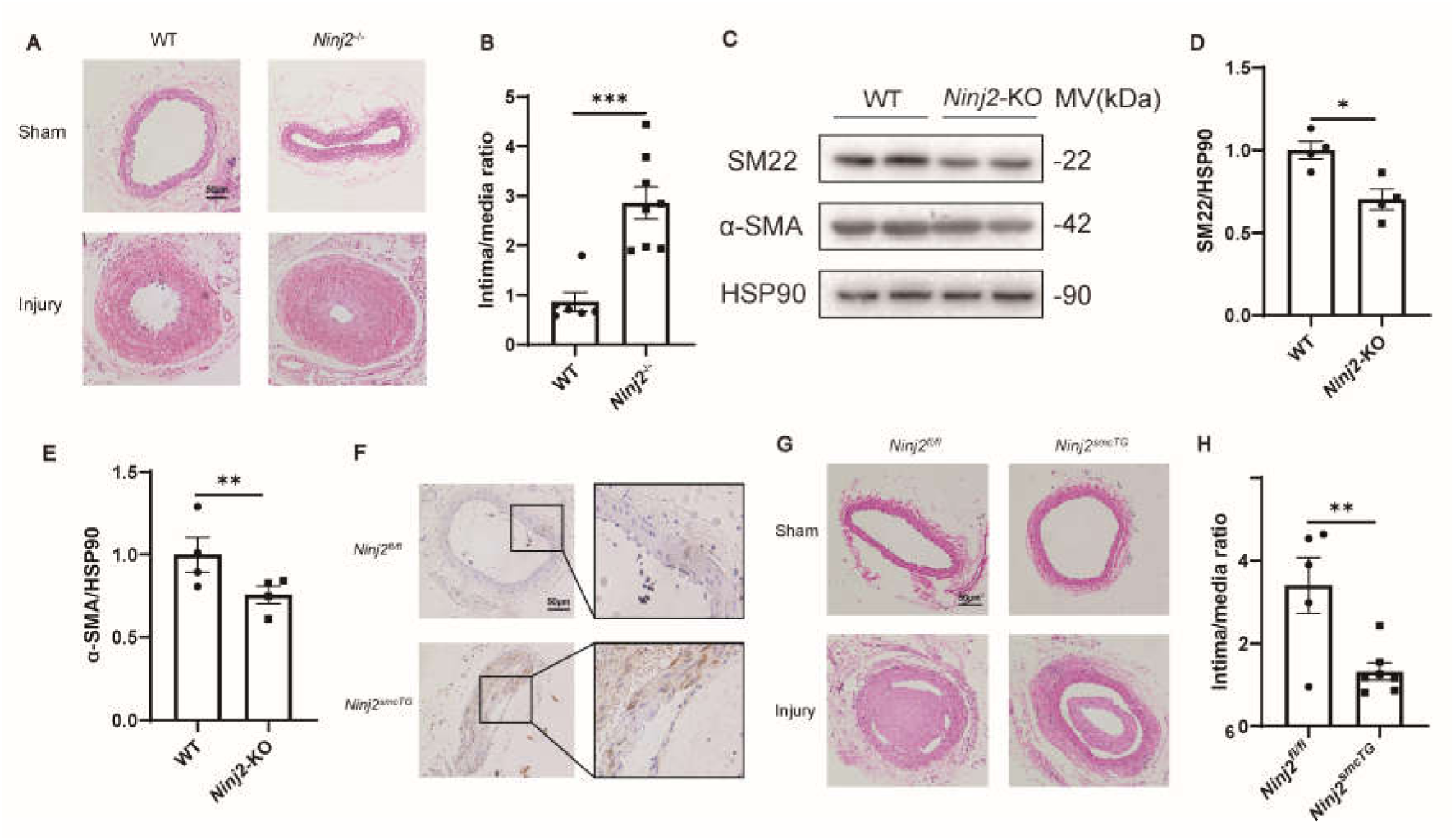
Ninjurin2 modulates neointimal hyperplasia and VSMC phenotypic plasticity following vascular injury in mouse model. A-B, Enhanced neointimal hyperplasia in *Ninj2*-deficient mice following carotid artery ligation. H&E staining of carotid arteries from 10-week-old *Ninj2^-/-^*knockout (KO) mice and wild-type (WT) littermates 28 days after ligation reveals significantly augmented neointimal formation in KO animals, as quantified by increased intima-media ratios compared with WT controls. Scale bar, 50 μm. n=6-8. C-E, Western blot analysis and the protein levels of contractile markers (α-SMA, SM22) in carotid arteries from Ninj2^-/-^ KO mice and WT controls 28 days after ligation (n=5/group). F, Immunohistochemical analysis of Ninjurin2 (brown signal indicating Ninjurin2 expression) demonstrates robust transgene expression in VSMCs of *Ninj2*^smcTG^ mouse mice compared to *Ninj2^fl/fl^*control group. Scale bar: 50 μm (20× magnification). G-H, Representative H&E-stained cross-sections of carotid arteries and quantitative analysis of intima/media thickness ratios from 10-week-old *Ninj2*^smcTG^ mouse and *Ninj2^fl/fl^* littermate controls 28 days after ligation surgery. Sham-operated vessels exhibited normal vascular architecture. Scale bars: 50 μm (20× magnification) (n=5-7). *p<0.01, **p<0.01, p was obtained using two-tailed Student’s t-test. Data represent mean ± SEM.

To ascertain the direct impact of Ninjurin2 in VSMCs phenotypic switching, we proceeded to investigate the effects of Ninjurin2 overexpression in neointimal formation using a VSMC specific overexpression of *Ninj2* (Mouse ortholog of Human *NINJ2* gene). We generated a knock-in mouse line, *Rosa26-CAG-Loxp-stop-loxp-Ninj2-IRES-EGFP*, where *Ninj2-IRES-EGFP* was integrated into the ROSA26 locus, flanked by loxP sites (*Ninj2^fl/fl^*). By crossing with *Tagln*-Cre transgenic mouse line, we produced VSMC-specific *Ninj2* overexpression mice (*Ninj2*^smcTG^). Immunostaining confirmed heightened Ninjurin2 expression in VSMCs *Ninj2*^smcTG^ mice when compared to control groups (Figure 1F). After 28 days of carotid artery ligation, H&E staining showed that a significantly reduced degree of neointimal formation in *Ninj2*^smcTG^ mouse compared with the *Ninj2^fl/fl^* control group (Figure 1G-H). These data suggest that *Ninj2* overexpression attenuates intimal hyperplasia subsequent to vascular injury.

Collectively, these results demonstrate a critical role for Ninjurin2 in mediating neointimal hyperplasia and modulating VSMC phenotypic plasticity following vascular injury.

### 3.2 Knockdown of Ninj2 expression in MOVAS promote VSMC phenotypic switching

Neointimal formation and intimal thickening are primarily attributed to the abnormal proliferation of VSMCs following their phenotypic switching and subsequent migration to the intima of blood vessels. To ascertain whether *Ninj2* regulates the phenotypic switching of VSMCs *in vitro*, we employed small interfering RNA (siRNA) to knock down *Ninj2* gene expression in mouse aortic smooth muscle cells (MOVAS). Subsequently, we assessed the expression of contractile protein markers using Western blot analysis. The results revealed a significant decrease in the expression levels of contractile markers, including SM22 and α-SMA, upon *Ninj2* knockdown (Figure 2A-C). These results suggest that *Ninj2* is important in modulating the phenotypic switching of VSMCs from a contractile to a synthetic phenotype *in vitro*.

**Figure 2.**
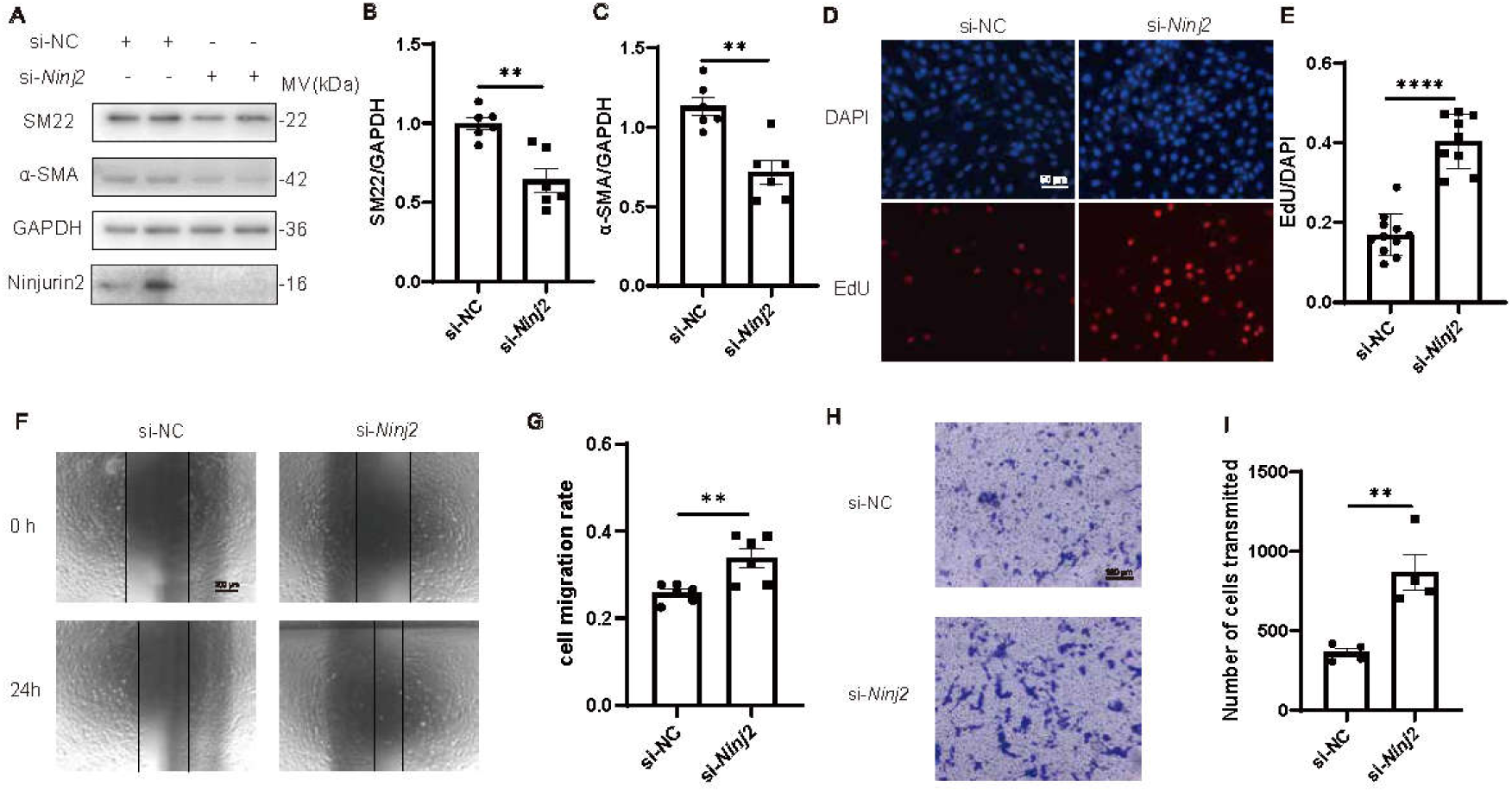
Ninj2 knockdown promotes phenotypic switching, proliferation, and migration in MOVAS vascular smooth muscle cells. A-C, Western blot analysis demonstrates decreased expression of contractile markers (SM22α, α-SMA) upon siRNA-mediated *Ninj2* knockdown in MOVAS, indicating enhanced phenotypic switching from contractile to synthetic states. Quantification of band intensities normalized to GAPDH is shown. **p<0.01. D-E, EdU proliferation assay reveals increased DNA synthesis (red fluorescence) in Ninj2 knockdown MOVAS at 36h post-transfection compared to si-NC controls. Scale bars: 50 μm. Quantification of EdU positive cells per field is shown (mean ± SEM, n=10 fields/group). ****p<0.0001. F-G, Scratch-wound assays confirm enhanced migratory capacity in *Ninj2*-knockdown MOVAS at 24h post-wounding. Dashed lines indicate initial wound margins. Migration distance was quantified as wound closure percentage (right panel; mean ± SEM, n=6 experiments). **p<0.01 vs. si-NC. H-I, Transwell migration assays further validate increased VSMC motility following *Ninj2* silencing (crystal violet-stained cells, lower chamber). Scale bars: 50 μm. Migrated cells were counted per high-power field (right panel; mean ± SEM, n=4 replicates). **p<0.01 vs. si-NC.

To determine whether *Ninj2* modulates the proliferation and migration of VSMCs, we conducted an EdU incorporation assay to assess cellular proliferation. Initially, we utilized siRNA to knockdown *Ninj2* expression in MOVAS. At 36 hours post-transfection, we evaluated the proliferation of MOVAS using an EdU incorporation assay. Figure 2D illustrates that the number of red fluorescent EdU-positive cells, indicative of proliferating cells, was markedly higher in MOVAS cells with *Ninj2* knockdown compared to the si-NC control group(Figure 2E). Furthermore, we observed that the knockdown of *Ninj2* in MOVAS cells significantly enhanced VSMC migration in both scratch-wound healing assays (Figure 2F-G) and Boyden-chamber Transwell assays (Figure 2H-I).

### 3.3 Ninj2 modulate PDGF-BB induced phenotypic switching, proliferation and migration of VSMC

Previous research has established that Ninjurin2 interacts with the PDGFR-β in glioma cells, and the primary natural ligand of PDGFR-β[11], PDGF-BB, is a critical factor in neointimal formation following vascular injury[19]. This is due to its potent stimulatory effects on VSMCs proliferation and migration through PDGFR-β[20]. Consequently, PDGF-BB is frequently utilized as a cellular model to investigate the signaling events implicated in neointimal formation. Building on this foundation, our study aimed to further investigate whether *Ninj2* modulates the phenotypic switching process of VSMCs under PDGF-BB stimulation.

As depicted in Figure 3A, the expression levels of contractile-type markers of VSMCs, including SM22 and α-SMA, were diminished in MOVAS cells upon stimulation with 20 and 40 ng/ml concentrations of PDGF-BB(Figure 3B-C). Furthermore, when *Ninj2* expression was knocked down by siRNA, the expression levels of SM22 and α-SMA were found to decrease even more significantly (Figure 3D-F). These findings suggest that the downregulation of *Ninj2* enhances the phenotypic switching of VSMCs from a contractile to a synthetic phenotype i*n vitro*, under the istimulation of PDGF-BB. Consistent with our previous findings, we utilized EdU incorporation assay(Figure 4A) and scratch-wound healing assays (Figure 4B) to demonstrate that proliferation and migration of VSMCs was markedly enhanced following *Ninj2* gene knockout upon stimulation with PDGF-BB.

**Figure 3.**
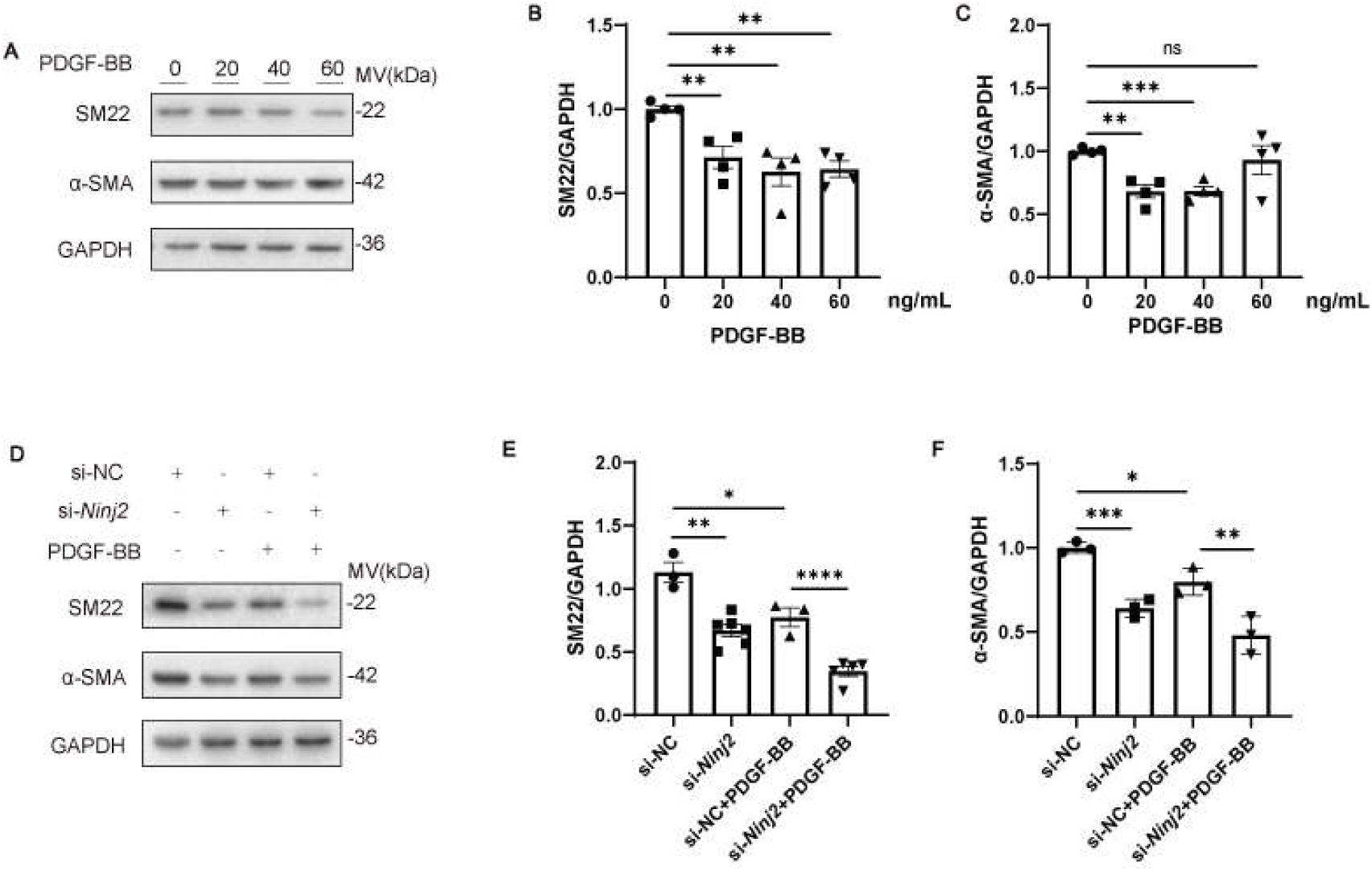
*Ninj2* modulates PDGF-BB-induced phenotypic switching of MOVAS vascular smooth muscle cells. A-C, PDGF-BB dose-dependently suppresses contractile phenotype markers in MOVAS. Representative immunoblots (A) and (B-C) quantitative analysis of SM22α and α-SMA expression in MOVAS treated with 0, 20, 40, or 60 ng/ml PDGF-BB for 24 hours. Data represent mean ± SEM (n=4 biological replicates per group). ***p<0.001, **p<0.01 vs. 0 ng/ml group by one-way ANOVA. D-F, *Ninj2* knockdown amplifies PDGF-BB-mediated phenotypic switching. Representative immunoblots (D) and (E-F) quantified densitometry of SM22α and α-SMA in siRNA-mediated *Ninj2*-knockdown MOVAS treated with 20 ng/ml PDGF-BB versus si-NC controls. Data shown as mean ± SEM (n=3-6 independent replicates). ***p<0.001, **p<0.01.

**Figure 4.**
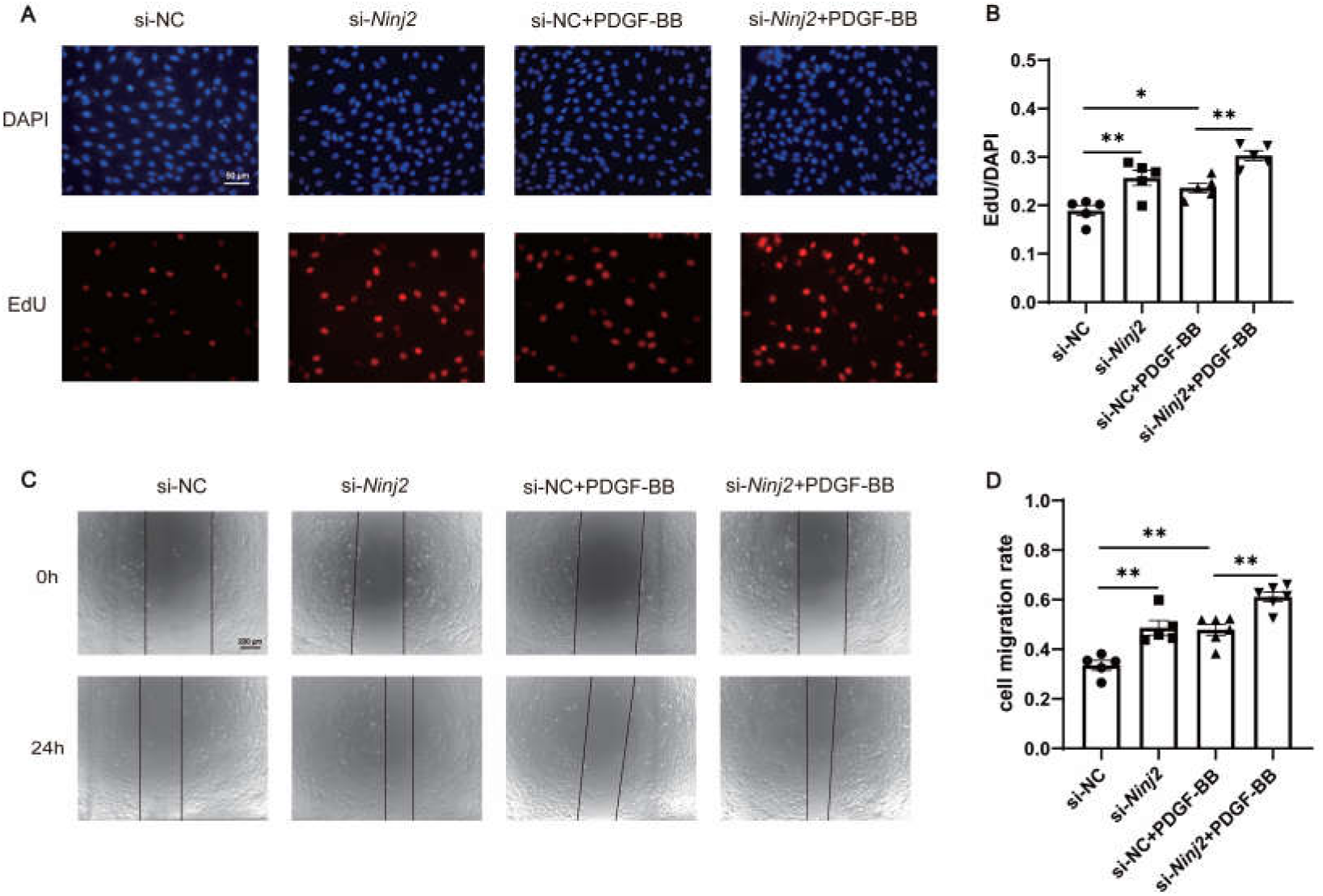
*Ninj2* modulates PDGF-BB-induced proliferation and migration of MOVAS vascular smooth muscle cells. A-B, EdU proliferation assay reveals increased DNA synthesis (red fluorescence) in Ninj2 knockdown MOVAS at 36h post-transfection compared to si-NC controls under 20ng/ml stimulation of PDGF-BB. Scale bars: 50 μm. Quantification of EdU positive cells per field is shown (mean ± SEM, n=5 fields/group). F-G, Scratch-wound assays confirm enhanced migratory capacity in *Ninj2*-knockdown MOVAS at 24h post-wounding under 20ng/ml stimulation of PDGF-BB. Dashed lines indicate initial wound margins. Migration distance was quantified as wound closure percentage (right panel; mean ± SEM, n=5 experiments). ***p<0.001, **p<0.01.

Collectively, these data suggest that the knockdown of *Ninj2* enhanced the PDGF-BB induced phenotypic switching, proliferation and migration of VSMCs.

### 3.4 The transmembrane domains of Ninjurin2 and the protein kinase domain of PDGFR-β is essential for interaction between Ninjurin2 and PDGFR-β

Previous studies have established that Ninjurin2 interacts with PDGFR-β in glioma cells. However, it remains unclear whether this interaction occurs in VSMCs and which key domain is essential for the interaction between Ninjurin2 and PDGFR-β. To elucidate the details of this interaction, our initial findings from immunofluorescence experiments conducted in MOVAS cells demonstrate that Ninjurin2 is predominantly co-localized with PDGFR-β on the plasma membrane, with partial co-localization observed in the cytoplasm (Figure 5A). Co-immunoprecipitation (Co-IP) showed that in HEK 293T cells with co-expression of GFP-PDGFR-β and FLAG- Ninjurin2, an anti-FLAG antibody successfully precipitated GFP-PDGFR-β (Figure 5B), and vice versa for the reciprocal Co-IP (Figure 5C).

**Figure 5.**
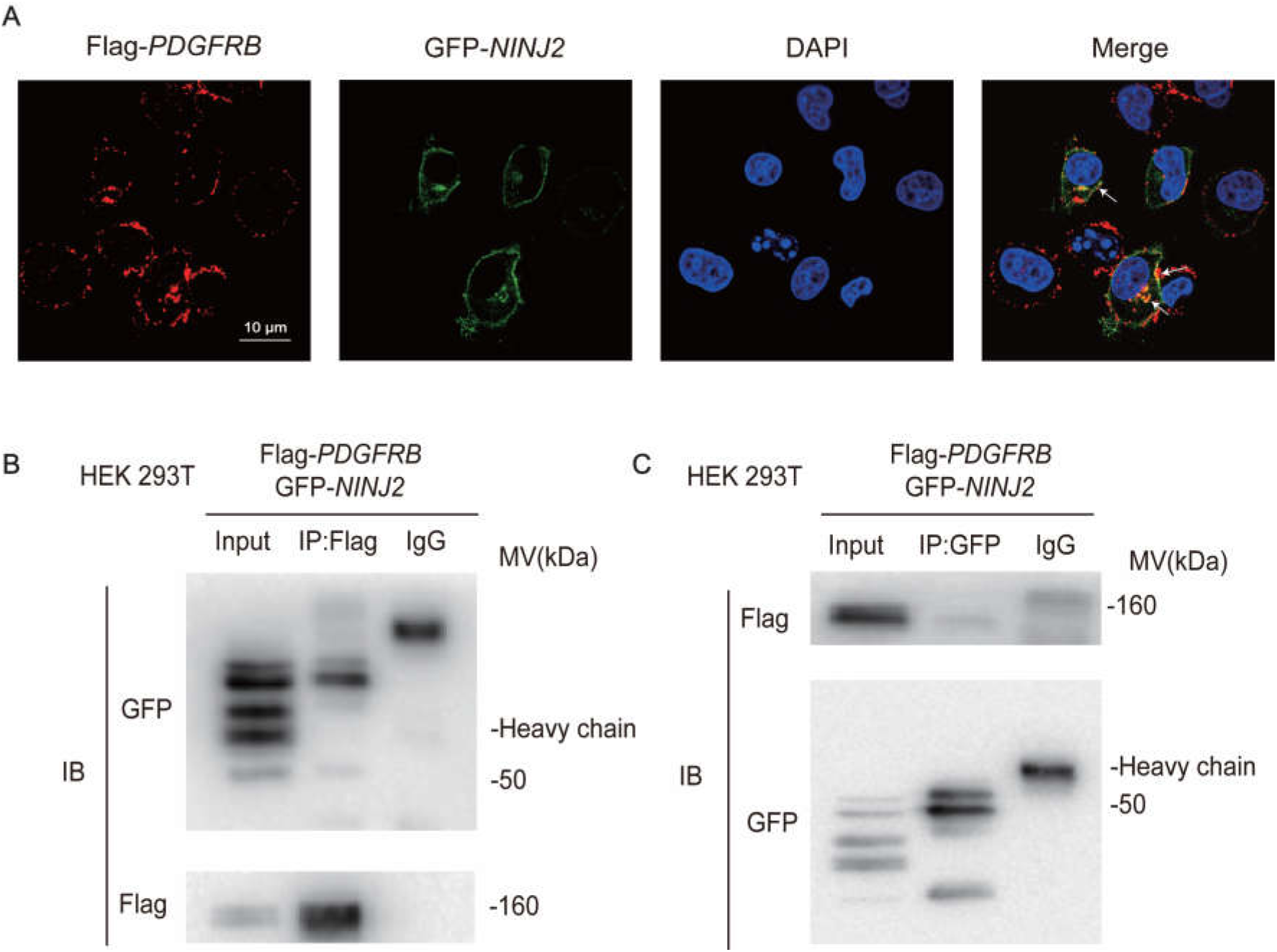
Co-localization and interaction between Ninjurin2 and PDGFR-β. A, Subcellular co-localization of Ninjurin2 and PDGFR-β in HeLa cells. Cells were co-transfected with GFP-tagged NINJ2 (pEGFP-N1-NINJ2; green) and Flag-tagged PDGFR-β (p3×Flag-CMV-10-PDGFRB; red). At 48 h post-transfection, cells were fixed, immunostained with anti-Flag primary and Alexa Fluor-conjugated secondary antibodies, and imaged by confocal microscopy. Yellow puncta in merged panels (arrows) indicate membrane- and cytoplasm-localized interaction sites. Scale bars: 10 μm. B–C, Co-immunoprecipitation (Co-IP) confirms direct Ninjurin2–PDGFR-β interaction. HEK293T cells were co-transfected with GFP-NINJ2 and Flag-PDGFR-β. B, Lysates precipitated with anti-Flag magnetic beads were immunoblotted (IB) using anti-GFP antibody. C, Reciprocal Co-IP: GFP-Trap magnetic bead pull-down followed by anti-Flag IB. IgG controls confirmed antibody specificity. Input lanes (5% total lysate) validate expression. IgG was used as a control antibody.

Based on the high resolution cryo-electron microscope atomic structure, Ninjurin2 contained two amphipathic helix domains in the N-terminal region (designated AH1 and AH2, encompassing amino acids 1 to 62), two transmembrane domains (labeled TM1 and TM2, spanning amino acids 63 to 97 and 98 to 125, respectively) and one short extracellular domain from the C-terminal region (126-142 amino acids). To ascertain the specific domains of Ninjurin2 responsible for interaction with PDGFR-β, we initially generated truncated plasmids expressing the AH1 and AH2 domains (amino acids 1-62) and the region encompassing TM1, TM2, and the C-terminal extracellular domain (amino acids 63-142) (Figure 6A). Co-IP experiments demonstrated that the full-length Ninjurin2 and the fragment containing amino acids 63-142 interacted with PDGFR-β, whereas the fragment spanning amino acids 1-62 did not (Figure 6B). Subsequent generation of serial truncated plasmids expressing TM1 (amino acids 63-97), TM2 (amino acids 98-125), and the C-terminal extracellular domain (amino acids 126-142) revealed that the TM1 and TM2 domains were capable of interacting with PDGFR-β (Figure 6C).

**Figure 6.**
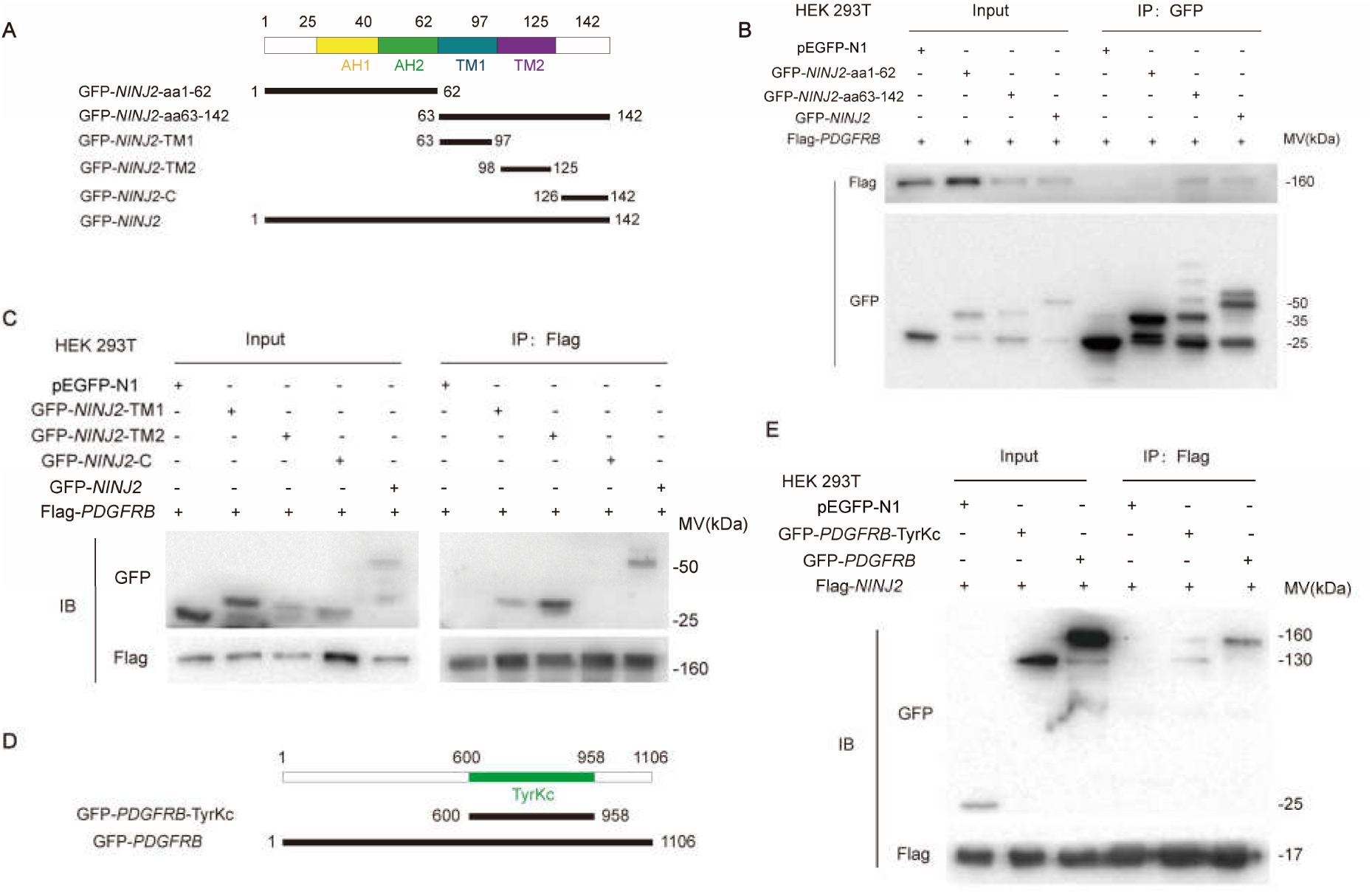
Domain mapping indicates the transmembrane domains of Ninjurin2 and the protein kinase domain of PDGFR-β is essential for interaction between Ninjurin2 and PDGFR-β. A, Domain architecture of Ninjurin2 and truncated constructs. Schematic depicts full-length Ninjurin2 (top) and deletion mutants used for interaction mapping (bottom), including GFP-NINJ2-aa1-62 contained N terminal region and two N-terminal amphipathic helices (AH1, residues 1–31; AH2, residues 32–62), GFP-NINJ2-aa63-142 contained two transmembrane domains (TM1, 63–97; TM2, 98–125) and C terminal region, GFP-NINJ2-TM1 contained transmembrane domain 1(63–97aa), GFP-NINJ2-TM2 contained transmembrane domain 1(98–125aa), GFP-NINJ2-C contained C terminal region(126-142aa). B, Co-IP analysis showing that GFP-NINJ2-aa63-142 which contained two transmembrane domains (TM1 and TM2) and C terminal region are necessary for PDGFR-β binding. HEK293T cells were cotransfected with PDGFR-β and full length or truncted Ninjurin2 expression plasmids (GFP-NINJ2, GFP-NINJ2-aa1-62,GFP-NINJ2-aa63-142) for 48 h, lysed, and used for immunoprecipitation. Anti-GFP was used for immunoprecipitation and anti-Flag was used for Western blotting. C, Further domain mapping indictaed TM1(63–97aa) and TM2(98–125aa) domains of Ninjurin2 were capable of interacting with PDGFR-β. D, Schematic of PDGFR-β domain architecture and the expression plamid architecture of the co-IP experiments. Truncted PDGFR-β contained tyrosine kinase domain domain (GFP-PDGFRB-TyrKc,600-958aa) and full length PDGFR-β were constructed. E, Co-IP analysis showing that the kinase domain (TyrKc) of PDGFRB interacts with Ninjurin2. HEK293T cells were cotransfected with Flag-Ninjurin2 and GFP tagged PDGFR-β or PDGFRB-TyrKc, Anti-Flag was used for immunoprecipitation and anti-GFP was used for Western blotting.

Ninjurin2 has previously been shown to interact directly with a variety of receptor tyrosine kinases (RTKs) in cancer cells, including epidermal growth factor receptor (EGFR), platelet-derived growth factor (PDGFR-β), and fibroblast growth factor receptor (FGFR). Our prior investigations also indicated that Ninjurin2 can directly interact with the insulin receptor (INSR)/insulin-like growth factor 1 receptor (IGF1R) in hepatocytes and adipocytes. All these Ninjurin2 interacted receptors have protein kinase domain, so we hypotheses that Ninjurin2 may interact with the protein kinase domain of PDGFR-β. Employing the SMART database, we predicted the protein kinase domain of PDGFR-β, which was found to span amino acids 600 to 958 (Figure 6D). We then constructed a truncated plasmid expressing the tyrosine kinase domain of PDGFRB (GFP-PDGFRB-TyrKc), and subsequent Co-IP experiments confirmed that the kinase domain (TyrKc) of PDGFRB interacts with Ninjurin2 (Figure 6E).

### 3.5 Ninjurin2 regulates focal adhesion kinase (FAK) pathway in VSMCs

To uncover the molecular mechanisms behind the phenotypic switching of vascular smooth muscle cells regulated by Ninjurin2, we conducted RNA-Seq analysis on MOVAS transfected with siRNA targeting *Ninj2* or a non-targeting control. Our findings revealed that the downregulation of *Ninj2* in VSMCs by siRNA led to the upregulation of 325 genes and the downregulation of 152 genes (with a|log2FoldChange| > 1.0). Gene Ontology (GO) differential expression analysis indicated that the differentially expressed genes were significantly enriched in functions such as developmental processes, extracellular matrix organization, cytoskeletal structure, multicellular organismal development, and regulation of molecular functions. KEGG pathway analysis identified several pathways that were significantly enriched, including vascular smooth muscle contraction, the PI3K-AKT signaling pathway, focal adhesion signaling pathway, Rap1 signaling pathway, extracellular matrix (ECM) receptor interaction, and the MAPK signaling pathway (Figure 7A).

**Figure 7.**
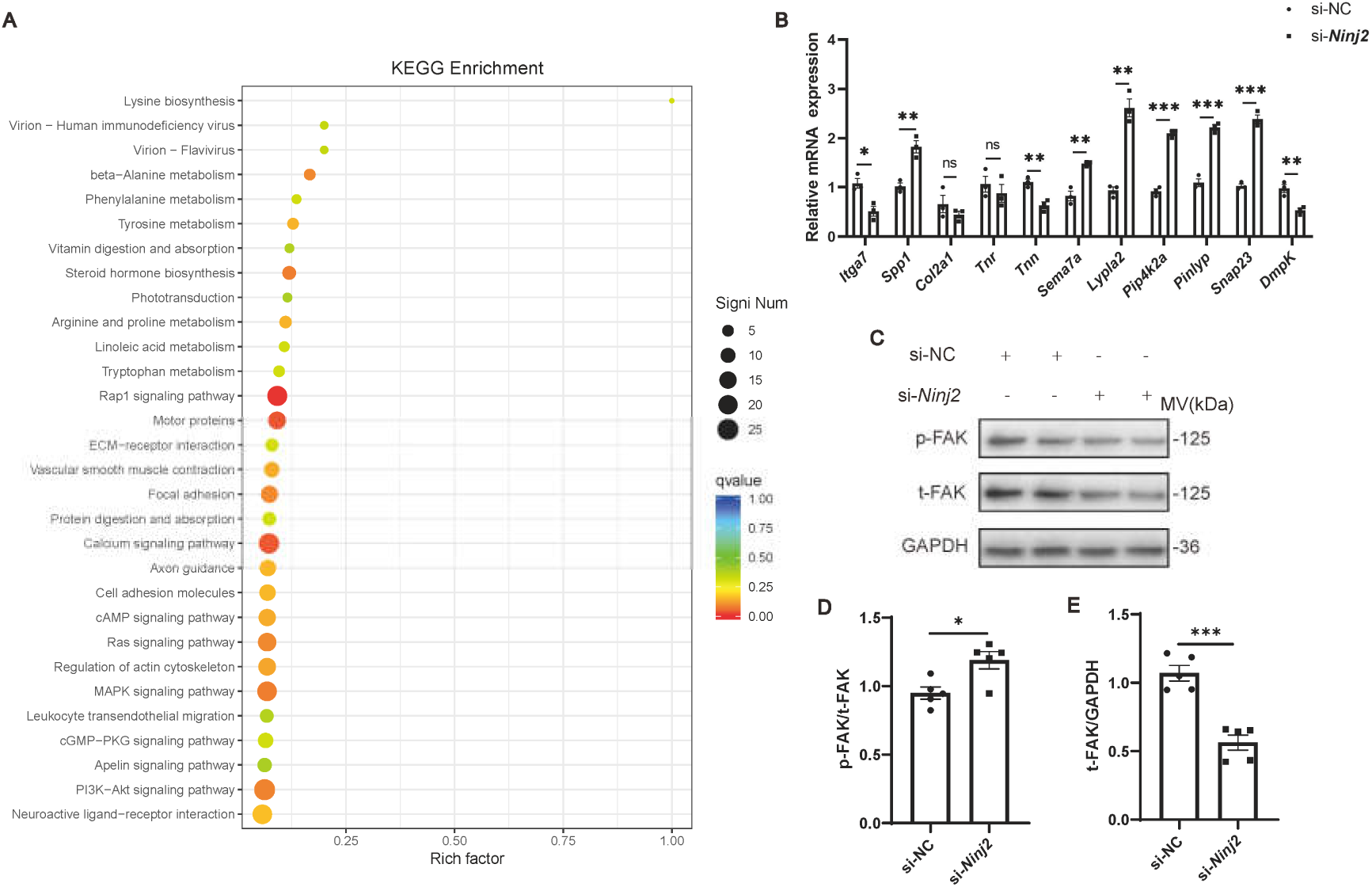
Transcriptomic profiling reveals Ninjurin2’s role in regulating VSMC phenotypic switching through FAK-mediated adhesion pathways. A, Differential gene expression and pathway analysis of RNA-seq data from MOVAS cells transfected with si-Ninj2 versus si-NC. Significantly enriched KEGG pathways among differentially expressed genes (DEGs; 325 upregulated, 152 downregulated. Thresholds: |log2FC| > 1.0, p < 0.05) (n = 3 biological replicates). Top pathways include ECM organization (GO:0030198), vascular smooth muscle contraction (hsa04270), and PI3K-AKT signaling (hsa04151). B, qRT-PCR validation of DEGs. Expression changes of select genes with TPM > 50, |log2FC| ≥ 1, and p < 0.05 were confirmed. *Itga7*, *Tnn*, and *Dmpk* were downregulated, while *Spp1*, *Sema7a*, *Lypla2*, *Pip4k2a*, *Pinlyp*, and Snap23 were upregulated. Data normalized to *TBP* (mean ± SEM, n = 3 replicated experiments; *p < 0.05,**p < 0.01, ***p < 0.001, two-tailed t-test). C-E, *Ninj2* regulates FAK protein phosphorylation and expression level. Western blot analysis and quantification of MOVAS cells transfected with si-Ninj2 or si-NC. Total FAK protein levels decreased upon Ninj2 knockdown.Phospho-FAK (Tyr397) levels increased, suggesting FAK activation. GAPDH served as loading control. Quantification shows band intensity ratios (p-FAK/total FAK, normalized to si-NC; mean ± SEM, n = 5).

In the present study, genes exhibiting a Transcripts Per Million (TPM) value exceeding 50, a significant |log2FoldChange| of at least 1, and a p-value of 0.05 or lower were chosen for validation via quantitative real-time fluorescence PCR (qRT-PCR). The qRT-PCR validation confirmed that the expression levels of Itga7, Tnn, and Dmpk were significantly downregulated following *Ninj*2 knockdown. Conversely, the expression of Spp1, Sema7a, Lypla2, Pip4k2a, Pinlyp, and Snap23 was significantly upregulated upon *Ninj2* knockdown (Figure 7B).

Intimal thickening due to neointima formation in atherosclerosis represents a vascular remodeling process, characterized by the migration and proliferation of VSMCs into the intima layer. Under normal physiological conditions, VSMCs are enveloped by the extracellular matrix (ECM), which includes collagen and adhesive proteins that interact with a variety of integrins. However, in pathological states, the ECM composition is altered due to enhanced secretion of matrix-remodeling enzymes [39]. Focal adhesion kinase (FAK) is pivotal in ECM and integrin signaling transduction [40, 41]. Typically, FAK is localized to the nuclei of VSMCs in healthy arteries, but cellular injury can trigger FAK activation and translocation [42]. The increased stiffness of the vascular wall during intimal hyperplasia activates FAK, thereby promoting VSMC proliferation and migration [43]. RT-qPCR data revealed that the knockdown of Ninj2 influences the expression of genes within the adhesion patch pathway, namely Spp1, Itga7, and Tnn, all of which are modulated by FAK. We further investigated whether *Ninj2* modulates downstream genes through the regulation of FAK expression using in vitro assays. MOVAS were transfected with siRNA targeting *Ninj2*, and subsequent Western Blot analysis was conducted to assess FAK protein expression and phosphorylation levels((Figure 7C-E). Relative to the negative control (NC) group, our study observed an increase in the phosphorylated FAK protein levels following *Ninj2* knockdown. Additionally, it was found that Ninj2 knockdown could decrease the overall FAK protein expression. Consequently, it is hypothesized that *Ninj2* may significantly contribute to VSMC proliferation and migration, potentially by regulating FAK signaling and impacting genes associated with the adhesion patch pathway, including Spp1, Tnn, and Itga7.

## 4. Discussion

Atherosclerosis, a chronic vascular disease, involves a critical phenotypic switching in VSMCs from contractile to synthetic states, which is central to its pathogenesis[21, 22]. While genetic variations in *NINJ2* are associated with risk of atherosclerosis and related conditions like ischemic stroke and CAD[23-25], the mechanisms of Ninjurin2’s impact on atherosclerosis, especially on VSMC phenotypic switching, are not fully understood. In the current study, we showed that ninjurin2 is a novel modulator for VSMC phenotypic switching and vascular remodeling. In vivo, *Ninj2* knockout (*Ninj2^-/-^*) mice exhibited increased neointimal formation post-vascular injury, associated with enhanced VSMC phenotype switching. In contrast, *Ninj2* specific overexpression in VSMCs of mice (*Ninj2*^smcTG^) attenuates intimal hyperplasia subsequent to vascular injury. In vitro knockdown of *Ninj2* in MOVAS cells promoted cell differentiation, proliferation, and migration. Finally, we demonstrated that ninjurin2 can interact directly with the protein kinase domain of PDGFRB, regulating the FAK signaling pathway— key regulators of VSMC phenotypic switching.

Compelling evidence suggests that VSMCs are instrumental in atherogenesis, predominantly contributing to the cellular composition of atherosclerotic lesions through phenotypic switching[26, 27]. Exposure to risk factors, including hypertension, lipid accumulation, inflammation, and infection, impairs vascular endothelial structure and function[28]. This impairment triggers the synthesis and release of vasoactive substances and cytokines, notably platelet-derived growth factor (PDGF)[29, 30]. Among its isoforms, PDGF-BB stands out as an early-identified promoter of phenotypic switching in VSMCs[31]. Acting on VSMC membrane receptors, predominantly PDGFRB, PDGF-BB activates a cascade of intracellular signal transduction pathways, inducing dedifferentiation in VSMCs[32]. Our research has uncovered a novel signaling pathway central to the pathobiology of neointimal formation following vascular injury, involving ninjurin2, PDGFRB, and the FAK signaling pathway (Figure 8). Through a series of deletion studies on Ninjurin2 and PDGFRB, we have pinpointed the TM1 and TM2 domains of Ninjurin2 as potentially critical in VSMC phenotypic switching, proliferation, and migration. Although cryo-electron microscopy has delineated the protein structure of Ninjurin2[33], the specific functions of each domain are not yet fully understood. Our investigation into the protein-protein interaction between Ninjurin2 and PDGFRB has not only clarified the significance of the TM1 and TM2 domains of ninjurin2 in VSMC phenotypic switching, proliferation, and migration, but also unveiled new avenues for understanding the structural and functional regulation of PDGFRB. Future studies will likely depend on the dissection of the structure and function of the Ninjurin2-PDGFRB protein complex to further elucidate the intricate mechanisms underlying their interaction and roles in vascular pathobiology.

**Figure 8.**
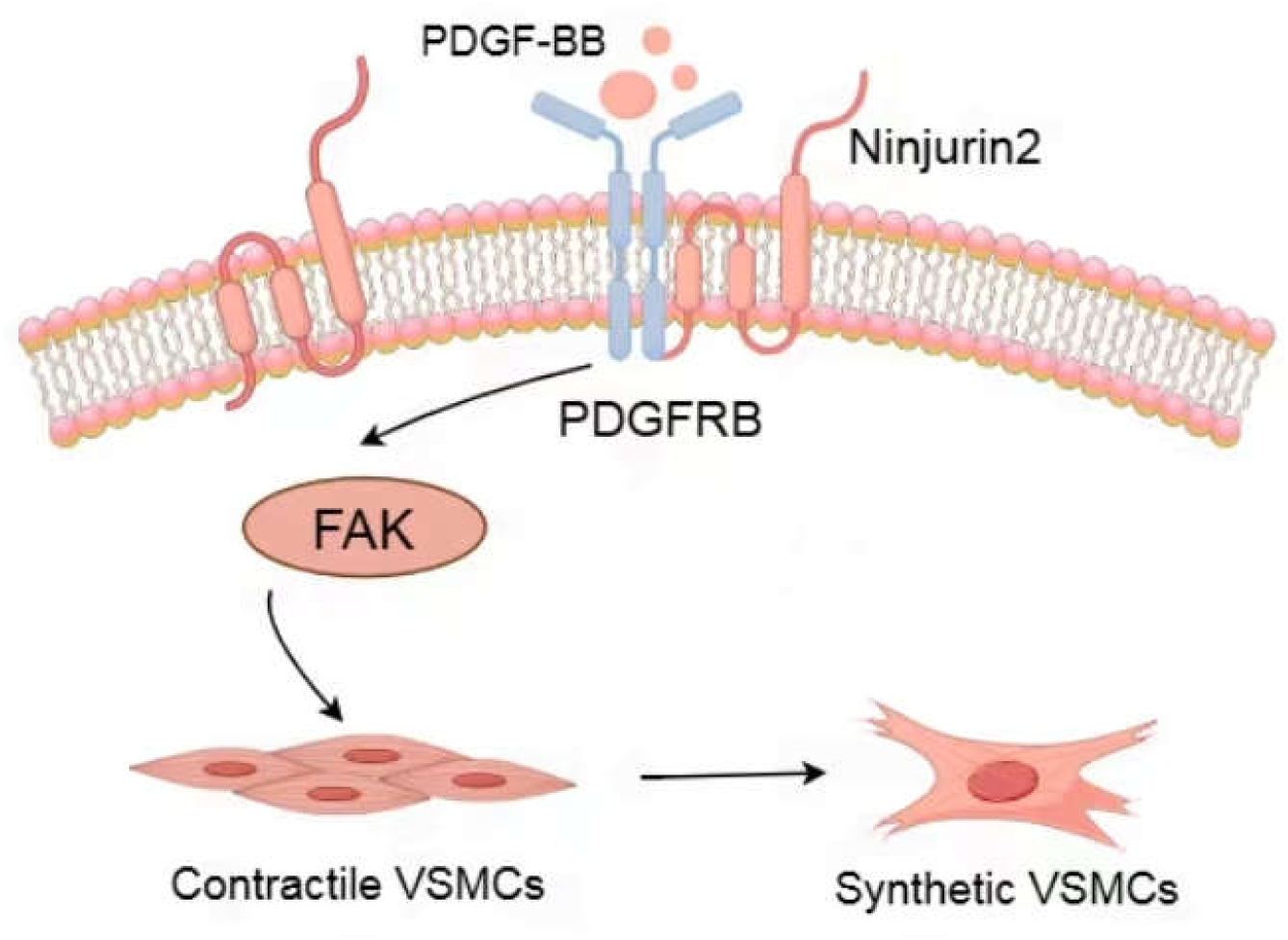
Mechanistic model of Ninjurin2-mediated regulation of VSMC phenotypic switching in vascular remodeling.

Our co-immunoprecipitation and domain-mapping experiments reveal that Ninjurin2 binds PDGFRB through its TM1/TM2 domains, while PDGFRB’s kinase domain mediates this interaction. This physical association likely modulates PDGFRB’s signaling output, as Ninjurin2 knockdown amplifies PDGF-BB-induced FAK phosphorylation and synthetic phenotype markers. Notably, FAK activation in VSMCs is a hallmark of ECM-integrin dysregulation during neointimal hyperplasia[34]. By suppressing FAK, Ninjurin2 may stabilize VSMCs in a quiescent state, counteracting PDGF-BB’s mitogenic effects. This aligns with transcriptomic data showing Ninjurin2 deficiency upregulates FAK-dependent genes (*Spp1*, *Sema7a*) and downregulates contractile markers (*Itga7*, *Tnn*). Further studies should determine whether Ninjurin2 directly inhibits PDGFRB kinase activity or sequesters it from downstream effectors.

In healthy blood vessels, smooth muscle cells maintain a contractile phenotype and express a range of contractile proteins, such as SM22 and α-SMA[35]. In response to vascular injury, contractile VSMCs down-regulate contractile protein expression and dedifferentiate into synthetic VSMCs, thus enhancing cell proliferation and migration, producing extracellular matrix and subsequently causing intimal hyperplasia[36]. Continuous dedifferentiation of VSMCs can lead to cardiovascular diseases, such as atherosclerosis, hypertension, restenosis after angioplasty, etc[37]. Especially in the process of atherosclerosis, phenotypic switching of VSMCs is an important step in vascular remodeling[38]. In this study, the expression levels of contractile proteins SM22 and α-SMA were decreased in both Ninj2 knockout mice and in vitro knockout mice compared to wild mice/controls. At the same time, the proliferation and migration ability of smooth muscle cells were increased after the low expression of Ninj2. In this study, it is concluded from both in vitro experiments and animal models that Ninj2 is involved in regulating phenotypic switching of VSMCs.

Ninjurin2 protein is a cell surface transmembrane protein that contains amphipathic helical regions AH1 and AH2 and two transmembrane domains TM1 and TM2 as shown by cryo-electron microscopy[33]. The protein family member Ninjurin1 protein can be cut into a soluble extracellular peptide in patients with coronary artery disease, which can regulate macrophage inflammation and improve atherosclerosis[39]. Subsequent studies have found that simulated short peptides containing specific amino acids of ninjurin1 can also ameliorate atherosclerosis[39]. This study shows that Ninjurin2’s transmembrane domains TM1 and TM2 can interact with PDGFRB proteins, which provides a new perspective for the study of Ninj2’s regulation of phenotypic switching of VSMCs. However, it is not clear whether the transmembrane domains TM1 and TM2 of Ninjurin2 protein are sufficient to function in place of full-length proteins, which may require further in vitro experiments or animal models to verify. Receptor tyrosine kinase family proteins have in common a conserved protein kinase domain, and when the signal is transmitted the protein forms a dimer on the membrane and activates its protein kinase function. In this study, GFP-PDGFRB-TyrKc truncated plasmid containing tyrosine kinase domain was constructed, and co-immunoprecipitation experiment proved that the interaction between PDGFRB protein and Ninjurin2 protein depends on the protein kinase domain of PDGFRB, which is not only beneficial to further explore the biological function of PDGFRB kinase domain. Moreover, it can provide a new idea for the study of targeted kinase inhibitors.

There is one limitation in this study, our study only investigated the molecular mechanism of Ninj2 mediating phenotypic switching of VSMCs at the animal level, but the function of Ninj2 in patients with atherosclerosis has not been studied.

## 5. Conclusions

In summary, this study found that Ninj2 gene knockout mice have serious intimal thickening in the presence of carotid artery injury, and Ninj2 VSMCs-specific overexpression mice can inhibit intimal hyperplasia after vascular injury, indicating that Ninj2 is involved in intimal lesions in the process of atherosclerosis. Studies at the cellular level have shown that knockdown of *Ninj2* results in the phenotypic switching from contractile to synthetic VSMCs, and abnormal proliferation and migration of VSMCs. Further studies have shown that down-regulation of *Ninj2* can increase phenotypic switching, cell proliferation and cell migration induced by PDGF-BB. This study further revealed that Ninj2 may regulate the intracellular FAK signaling pathway by interacting with PDGFRB. Subsequently, in this study, wer provide a new mechanism regulating phenotypic switching of VSMCs, and a new target for the treatment and prognosis of atherosclerosis.

## Author Contributions

Conceptualization, X.C. and W.P.; methodology, J.M., J.Z., X.Z.,T.L, H.Z., K.X.; investigation, X.X.; resources, J.M., J.Z., X.Z.; writing—original draft preparation, J.M., J.Z., X.Z.; writing—review and editing, X.C. and W.P; supervision X.C.; project administration, X.C.; funding acquisition, X.C. and W.P. All authors have read and agreed to the published version of the manuscript.

## Institutional Review Board Statement

Animal care and procedures were reviewed and approved by the Institutional Committee on Animal Care and Use, School of Medicine, Huazhong University of Science and Technology (No. [2016] ICUC469), and were conducted in compliance with institutional and relevant guidelines.

## Data Availability Statement

The original contributions presented in this study are included in the article/supplementary material. Further inquiries can be directed to the corresponding author.

## Acknowledgments

We acknowledge the support of the instrument share platform of collage of life science and technology of Huazhong University.

## Abbreviations

The following abbreviations are used in this manuscript:

VSMC: Vascular smooth muscle cell
Ninj2⁻/⁻: Ninjurin2 knockout mice
Ninj2ˢᵐᶜᵀᴳ: Smooth muscle-specific Ninjurin2 overexpressing mice
PDGFR-β/PDGFRB: Platelet-derived growth factor receptor-beta
FAK: Focal adhesion kinase
CAL: Carotid artery ligation
MOVAS: Mouse aortic smooth muscle cells
qRT-PCR: Quantitative reverse-transcription polymerase chain reaction
Co-IP: Co-immunoprecipitation
SNP: Single nucleotide polymorphism
CHARGE: Cohorts for Heart and Aging in Genomic Epidemiology
CAD: Coronary artery disease
EdU: 5-Ethynyl-2’-deoxyuridine
ECM: Extracellular matrix
GWAS: Genome-wide association study
TLR4: Toll-like receptor 4
AH1/AH2: Amphipathic helix domains 1 and 2
TM1/TM2: Transmembrane domains 1 and 2
TyrKc: Tyrosine kinase domain

